# Fine scale sampling unveils diazotroph patchiness in the South Pacific Ocean

**DOI:** 10.1101/2020.10.02.323808

**Authors:** Mar Benavides, Louis Conradt, Sophie Bonnet, Ilana Berman-Frank, Stéphanie Barrillon, Anne Petrenko, Andrea M. Doglioli

## Abstract

Diazotrophs are important contributors to reactive nitrogen availability in the ocean. Oceanographic cruise data accumulated along decades has revealed a heterogeneous distribution of diazotroph species at regional to global scales. However, the role of dynamic fine scale structures in distributing diazotrophs is not well understood. This is due to typical insufficient spatiotemporal resolution sampling and the lack of detailed physical studies in parallel. Here we show the distribution of five groups of diazotrophs in the South Pacific at an unprecedented resolution of 7-16 km. We find a patchy distribution of diazotrophs, with each group being differently affected by parameters describing fine scale structures. The observed variability could not have been revealed with a lower resolution sampling, highlighting the need to consider fine scale physics to resolve the distribution of diazotrophs in the ocean.

## Main body of text

The surface ocean is constantly stirred by currents that swirl and mix different seawater masses creating a dynamic mosaic of biogeochemical properties [1]. Numerical modeling and satellite data have shown that ‘fine scale’ structures such as filaments and eddies (with typical spatiotemporal scales of 1 to 100 km and days to weeks) impact the distribution of phytoplankton, primary productivity and carbon export in the ocean [2–4]. While these remote approaches provide valuable synoptic views at large scales, in situ sampling remains imperative to resolve the diversity, metabolism and trophic interactions of marine microbes at fine scales. However, the spatiotemporal resolution of typical at-sea sampling efforts is too coarse to resolve fine scale processes [5], which hampers our understanding of physical-biological interactions and their impact on marine microbes [1].

Understanding the effect of fine scales on biogeochemically relevant microbial groups is of particular importance. Dinitrogen (N_2_) fixers or ‘diazotrophs’ provide a significant source of bioavailable nitrogen in the ocean [6]. Some studies have shown a preferential accumulation of diazotrophs in anticyclonic eddies [7, 8], where eddy pumping deepens isopycnals impoverishing surface waters in inorganic nitrogen presumably favoring diazotroph growth [reviewed in 9]. However, other studies have reported accumulations of diazotrophs in cyclonic eddies instead, attributed to wind-driven Ekman pumping [10]. Diazotrophs encompass a wide diversity of prokaryotic microorganisms with different tolerance to environmental and nutrient availability conditions. For example, cyanobacteria such as *Trichodesmium* and UCYN-B abound in oligotrophic (sub)tropical waters, while UCYN-A displays a wide geographical distribution spanning from the tropics to the polar seas [6]. Non-cyanobacterial diazotrophs cannot photosynthesize and obtain carbon and energy from organic matter compounds, which presents fundamentally different bottom-up controls as compared to cyanobacterial diazotrophs [11]. With such divergent physiologies, it is unlikely that different diazotrophs respond to fine scale physical forcing in the same way.

Resolving these ambiguities needs coupling fine scale structure and diazotroph activity/abundance measurements at high spatiotemporal resolution. The vast majority of diazotroph activity/abundance data available in the literature up to 2012 stemmed from locations situated ~160 km apart (median distance between stations in the database compiled by Luo et al [12]). Recent methodological developments have provided underway N_2_ fixation and diazotroph abundance data at a spatial resolution of ~18 km [13, 14]. While such resolution may be appropriate to resolve fine scale processes, previous studies have not sought to explain diazotroph distributions through the lens of fine scale oceanography.

Here we investigate the fine scale variability of diazotrophs in the Southwest Pacific Ocean during a cruise in austral summer 2019 (Fig. 1). Zones of intense fine scale activity were selected for high-resolution sampling according to satellite and Lagrangian product maps received onboard on a daily basis [15]. These included sea surface temperature, chlorophyll, geostrophic current based on absolute dynamic topography (ADT), and the Lagrangian products finite size Lyapunov exponents (FSLE) and Okubo-Weiss parameter (OW) (Supplementary Information; Table S1; Fig. S1). Planktonic biomass was collected with an automated filtration system at an unprecedented resolution of 7-16 km. DNA extracted from the filters was used to quantify five diazotroph groups (*Trichodesmium*, UCYN-A1, UCYN-B, UCYN-C and Gamma A) in quantitative PCR assays (Supplementary Information). Only in zone 3 inorganic nutrient concentrations and N_2_ fixation rates were also measured (Supplementary Information).

**Fig. 1:**
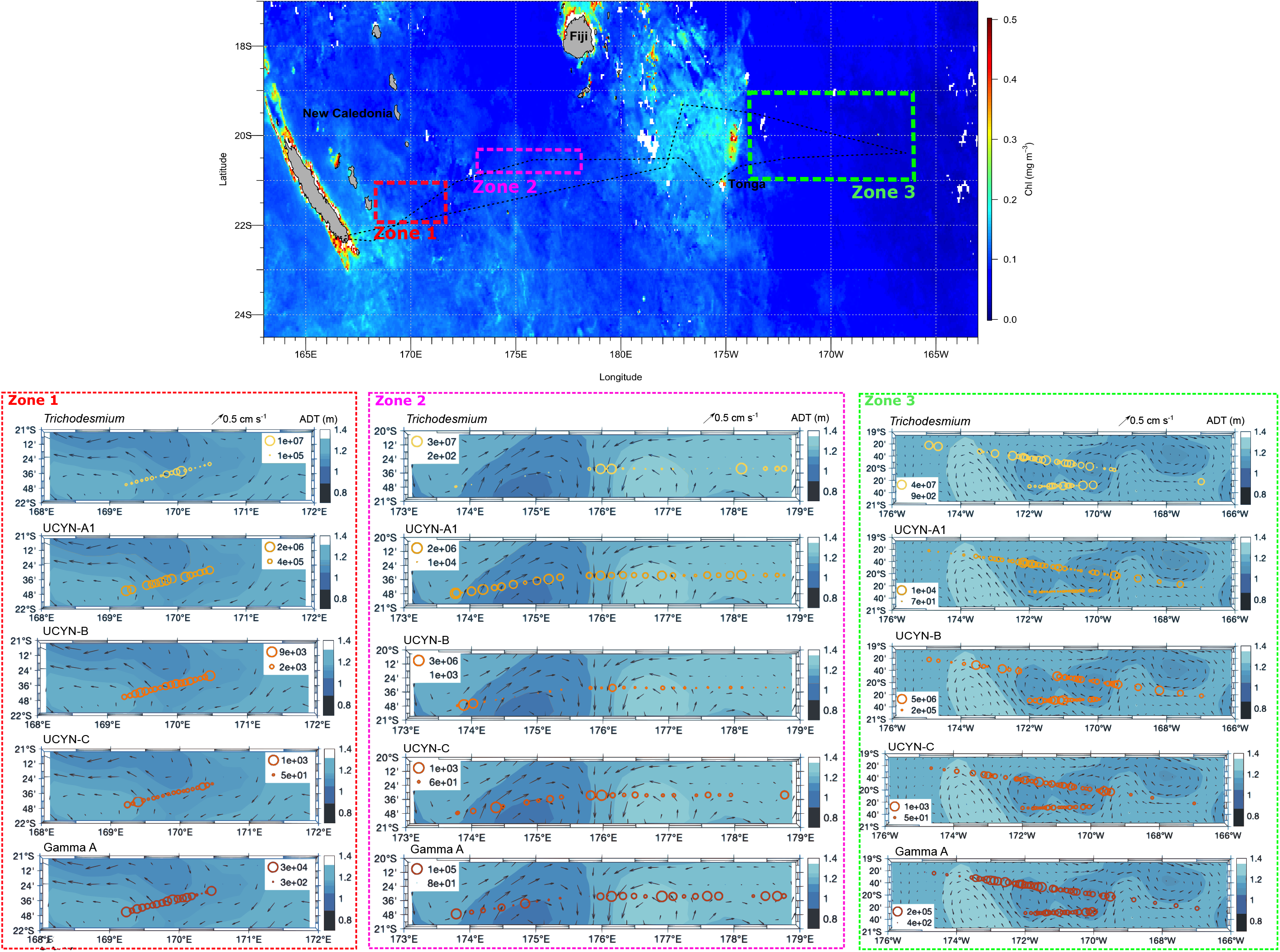
Fine scale resolution abundances of diazotrophs along three selected sampling zones. The central image shows a chlorophyll MODIS composite averaged for November 2019 at a resolution of 4 km and the location of the three selected sampling zones. Zoom-out panels (in dotted squares) show the abundance of diazotrophs in each selected zone as *nifH* gene copies per liter of seawater. Diazotroph abundances (*nifH* gene copies L^−1^) are superimposed on absolute dynamic topography (ADT, color scale) and geostrophic velocity (arrows). ADT data were retrieved for each zone on 2^nd^, 4^th^ and 22^nd^ November 2019 for zones 1, 2 and 3, respectively. To provide an overview, this figure only presents abundances of *Trichodesmium*, UCYN-A1 and Gamma A. The abundances of the other groups (UCYN-B and UCYN-C) are shown in Fig. S3.

Our results reveal a patchy distribution of diazotrophs, driven by a heterogeneous effect of fine scale parameters on each diazotroph group (Fig. 1; Fig. S2; Table S2). *Trichodesmium* correlated positively with ADT (Fig. 1; Fig. S2; Table S2), and accumulated at positive-negative OW transition and high FSLE regions located at ~170°E in zone 1, 176°E in zone 2 and 171°W in zone 3 (Fig. 2). High FSLE values depict fronts created by horizontal transport, which are susceptible to accumulate floating particles such as *Trichodesmium* colonies which harbor gas vesicles [16]. The accumulation of *Trichodesmium* at the convergence of two counter-rotating eddies in zone 2 (maximum abundance of 3 ×10^7^ *nifH* gene copies L^−1^; Fig. 1) was likely driven by Langmuir circulation [17]. Remarkably, the temperature of the two counter-rotating eddies of zone 2 differed by >1.5°C (Fig. S4). The abundance of UCYN-A1 ranged between 10^4^ and 10^6^ *nifH* gene copies L^−1^, with lower abundances along the southern transect in zone 3 (Fig. 1). UCYN-A1 were more homogeneously distributed independently of fine scale features and were negatively related to FSLE (Table S2) and to *Trichodesmium* (Figs. S2, S3), adding up to the antagonistic biogeographic trends of these two diazotrophs groups witnessed at larger spatial scales [12]. UCYN-B were more abundant in zone 3 (up to 10^6^ *nifH* gene copies L^−1^) than in zones 1 and 2 (up to 10^3^ *nifH* gene copies L^−1^) (Fig. 1). They were relatively more abundant at the edges of eddies (Fig. 1) coinciding with higher FSLE values (Fig. 2), although they were not statistically related to physical parameters (Table S2). UCYN-C were significantly related to both ADT and FSLE (Table S2), but were the least abundant group (up to 10^3^ *nifH* gene copies L^−1^; Fig. 1) likely due to their presumed coastal origin [6]. Finally, Gamma A were significantly related to ADT (Table 2) and accumulated in frontal zones with high FSLE (up to 10^5^ *nifH* gene copies L^−1^), which agrees with their putative particle-attached lifestyle [18].

**Fig. 2:**
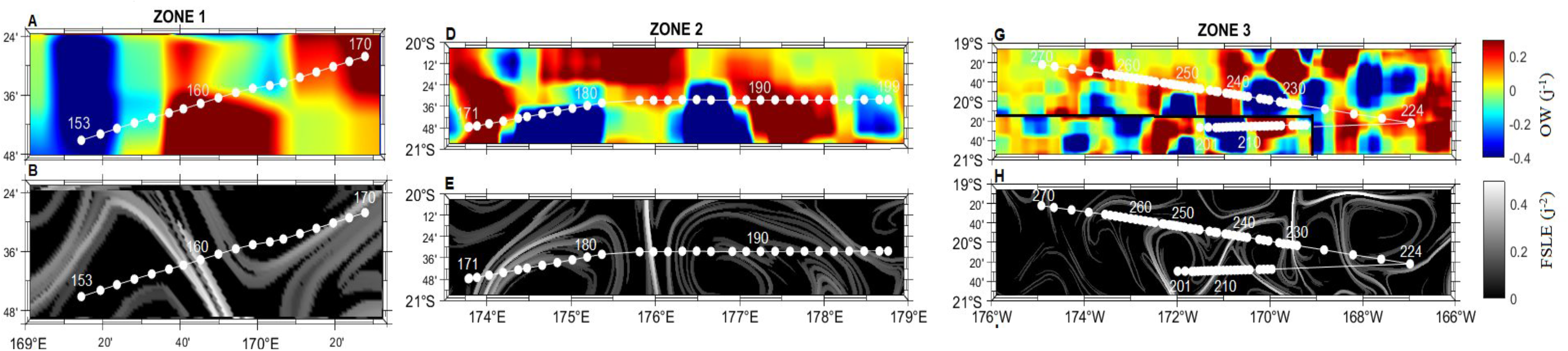
Lagrangian diagnostics parameters. Panels a), d) and g) show values of the Okubo-Weiss (OW) in zones 1, 2 and 3, respectively. Panels b), e) and h) show values of finite Lyapunov finite size Lyapunov exponents (FSLE) in zones 1, 2 and 3, respectively. White dots represent sampling locations.

In zone 3, nitrate concentrations were typically <0.2 μM (Fig. S5a). The relatively high concentrations of phosphate east of ~175°W (Fig. S5b) likely sustained only moderate N_2_ fixation rates (1-5 nmol N L^−1^ d^−1^; Fig. S5c), which correlated with the abundance of *Trichodesmium* (Spearman, p= 0.008). Low N_2_ fixation rates in this region are thought to be caused by the scarcity of iron east of the Tonga volcanic arc [19]. Nutrients and temperature typically used to define the biogeography of diazotrophs on regional to global scales [12].

Despite the remarkably homogeneous distribution of these factors in zone 3 (Fig. S4g, Fig. S5a-b), the distribution of diazotrophs revealed high spatial variability at the fine scale (Fig. 1). Such variability would have gone unseen at a coarser resolution, stressing the role of fine scale dynamics in diazotroph distribution. The patchiness observed likely responds to a combination of bottom-up and top-down interactions between diazotrophs’ competitors, predators and the surrounding biogeochemical environment. Documenting such mechanisms at fine scale resolution warrants exciting research avenues in the near future.

## Acknowledgements

This study was funded by the projects TONGA (ANR-18-CE01-0016, LEFE-CyBER, Fondation A-Midex, Flotte océanographique française) to S. Bonnet and C. Guieu, DEFINE (LEFE-CyBER) to Mar Benavides), and OASIS (Thomas Jefferson Fund) to S. T. Wilson and M. Benavides). The development of SPASSO is supported by TOSCA/CNES and by EU program Copernicus Academy. L. Conradt was funded by a Master degree internship from TONGA A-MIDEX. The authors would like to thank the crew and technical staff of R/V *L’Atalante* as well as the scientists that participated in sample acquisition and equipment installation onboard (C. Lory, K. Sellegri and F. Gazeau), and sample analyses in the lab (S. Nunige, O. Grosso, A. Torremocha). The MODIS chlorophyll image used in Fig. 1 is a courtesy of J. Uitz. The authors greatly acknowledge the OMICS platform at the Mediterranean Institute of Oceanography (Marseille, France) for access to their facilities.

