## Supplementary material for "Fine scale sampling unveils diazotroph patchiness in the South Pacific Ocean": DEFINE_SI.pdf

### Supplementary Information

#### Table of contents

1. Supplementary Methods
2. Supplementary Tables
3. Supplementary Figures
4. Supplementary References

### 1. Supplementary Methods

*Fine scale features and high-resolution biomass sampling*

The TONGA cruise ("shallow hydroThermal sOurces of trace elements: potential impacts on biological productivity and the bioloGicAl carbon pump"; http://tonga-project.org, doi: 10.17600/18000884) took place onboard the *R/V L'Atalante* between 31$^{st}$ October and 6$^{th}$ December 2019. High-resolution sampling while transiting between stations targeted fine scale structures shown in satellite product reports sent by the land team to embarked scientists on a daily basis. The daily report on fine scale variability was processed with the software 'Satellite-based Sampling for Oceanographic cruises' (SPASSO: 'Software Package for an Adaptive Satellite-based Sampling for Oceanographic cruises' https://spasso.mio.osupytheas.fr/; [2]), which we have previously used to study the fine scale distribution of different planktonic groups in the Mediterranean Sea, Southern and South Pacific Oceans [3–6]. SPASSO reports included images of sea surface temperature (SST), chlorophyll and geostrophic current based on absolute dynamic topography (ADT), obtained from the Copernicus Marine Environment Monitoring Service (https://marine.copernicus.eu/). The SPASSO products provided also included finite size Lyapunov exponents (FSLE) integrated over 30 days, and the daily values of the Okubo-Weiss parameter (OW). FSLE values indicate the exponential rate of particle trajectory separation, allowing to identify Lagrangian Coherent Structures (LCS) from altimetry data as candidates for fine scale frontal regions [1]. The OW parameter compares the importance of deformation of the current velocity field with respect to its rotation. Negative values of OW quantify the strength of an eddy, being its velocity field dominated by rotation.

SPASSO images were used to select three zones of relevant fine scale activity where to sample in high-resolution. Zones 1 and 2 ranged between 169.24°E - 170.46°E and between 173.78°E - 178.75°E, respectively. Zone 3 included one west-to-east (forward) and one east-to-west (return) transect, which ranged between 171.99°W - 169.95°W and 166.98°W - 174.92°W, respectively. Zone 1 was sampled on 2nd November and included 18 stations separated by an average distance of 4 nm. Zone 2 was sampled between 4th and 5th November and included 30 stations with an average separation of 8 nm. Finally, zone 3 included 23 stations on the forward transect sampled on 17th November with an average separation of 4 nm, while the return transect included 47 stations separated by 9 nm on average.

In each zone seawater was pumped from 5 m depth through the ship's underway outlet, which was washed with concentrated bleach and flushed with demineralized water for several hours before the cruise. Temperature and salinity were measured with a SBE21 thermosalinograph (Sea-Bird Scientific, Bellevue, WA, USA) and binned at 1 min. Fluorescence data was recorded every 10-15 min with a Wetstar fluorimeter (Sea-Bird Scientific). Plankton biomass was collected by an OCE-5 autosampler (Oceomics, Fuerteventura, Spain) connected to the ship's underway. Briefly, the OCE-5 redirected seawater through six ports (similar to the device described in [7]). Each port was fitted with 25 mm diameter 0.2 µm pore size polysulfone filters (Supor, Pall Gelman, Port Washington, NY, USA). The OCE-5 was programmed to filter one sample every 30 min when sailing over target zones. The tubing and connections of the OCE-5 were thoroughly flushed with 70%

EtOH and Milli-Q water between sampling zones. Filtered volumes ranged between 1200 and 2000 ml depending on biomass loading, with filtration times ranging from 10 to 22 min. Filters were stored in bead beater tubes containing a mix of 0.1 mm and 0.5 mm diameter glass beads (Oakton Instruments, Vernon Hills, IL, USA) and immediately stored at -80ºC.

*DNA extractions and quantitative PCR (qPCR)*

DNA was extracted using the DNeasy Plant Mini Kit (Qiagen, Courtaboeuf, France) with additional freeze-thaw bead beating and proteinase K steps before the kit purification and elution to 80 µl in RNase-free water as previously described [8].

The abundance of diazotrophs was determined using TaqMan qPCR assays and previously published primer-probe sets for *Trichodesmium,* UCYN-A1, UCYN-B, UCYN-C and γ-24474A11 (the latter hereafter referred to as Gamma A) [8–10]. The qPCR was run in 25 µl reactions consisting of 12.25 µl TaqMan PCR Master Mix (Applied Biosystems, Villebon Sur Yvette, France), 1 µl of the forward and reverse primers at 10 µM (HPLC purified, Eurofins, Nantes, France), 0.25 µl probe at 10 µM, 8.25 µl PCR grade water, 0.25 µl bovine serum albumin at 10.08 µg $\mu l^{-1}$, and 2 µl standard or template sample. The qPCR program was run on a CFX96 Real-Time System thermal cycler (BioRad, Marnes-la-Coquette, France) and consisted of 2 min at 50ºC, 10 min at 95ºC continued by 45 cycles of 15 s at 95ºC and 1 min at 64ºC. The annealing temperature was changed to 60ºC for UCYN-A1 qPCR runs [10]. Standard dilutions ($10^7$-$10^1$ gene copies) were run in duplicate, and samples and no-template controls (NTCs) in triplicate. NTCs did not show any amplification. The efficiency was 98 - 113%. Inhibition tests were carried out on all samples and each primer-probe set by adding the 2 µl of the $10^5$ copy standard to each sample. No inhibition was observed. The limit of detection and detected but not quantifiable limits were 1 and 8 gene copies per reaction, respectively.

*Nutrients and $N_2$ fixation rates*

Only on zone 3, samples were collected manually from the underway outlet for dissolved inorganic nutrient and $N_2$ fixation rate analyses. For nutrients, seawater was collected in acid-clean 20 ml polyethylene tubes and stored at -20ºC until analyses. Ashore, samples were analyzed after Aminot and Kérouel [11] on a Bran Luebbe AA3 autoanalyzer. Detection limits were 0.05 µM for both nitrate and phosphate. $N_2$ fixation rates were measured from 2.3 l samples collected on transparent polycarbonate bottles and amended with 2 ml 98.9 atom % $^{15}N_2$ gas (Euroiso-top, Saint-Aubin, France). Bottles were inverted 30 times and incubated in surface water circulated on-deck incubators for 24 h. Incubations were terminated by filtration onto precombusted 25 mm GF/F filters (GE Healthcare, Little Chalfont, UK) and analyzed on an Integra CN elemental analyzer coupled to an isotope ratio mass spectrometer (Sercon Instruments, Crewe, UK) as previously described [12]. To avoid an underestimation of $N_2$ fixation rates due to the incomplete dissolution of $^{15}N_2$ gas into seawater samples [13], the $^{15}N$ atom % enrichment of incubated seawater was measured from 12 ml headspace-free samples collected in exetainer tubes (Labco, Ceredigion, UK), stored at 4ºC and analyzed on a membrane inlet mass spectrometer (Pfeiffer, MD, USA) after Kana et al. [14]. The high-resolution sampling approach applied here did not permit sampling in triplicates for $N_2$ fixation rate measurements using $^{15}N_2$ isotope labeling. Hence, minimum quantifiable rates (MQR) are not reported. Nevertheless, companion measurements of $N_2$ fixation rates during the same cruise, which were labeled with the same volume of $^{15}N_2$ gas, equally incubated and analyzed with the same EA-IRMS and MIMS equipment.

*Statistical analyses*

Outlier values of diazotroph *nifH* gene copy abundance were removed by the interquartile range method. *NifH* gene copy number data were normally distributed and hence Pearson correlations were used to test their correlation with environmental data (i.e. temperature, salinity and fluorescence) using the Hmisc and corrgram packages in R [15, 16]. Fine scale structure parameters included FSLE, OW and ADT (see above).

An redundancy analysis (RDA) triplot was computed on Hellinger-transformed *nifH* gene copy abundance data as response variables and fine scale structure data as explanatory variables using the R package vegan [17]. To avoid misinterpreting zero values of FSLE as absence of data in RDA calculations, FSLE was treated as a non-quantitative variable with two levels: i) FSLE>0.05 $d^{-2}$ indicating the presence of Lyapunov coherent structures (LCS) such as fronts or filaments with an intensity corresponding to the FSLE value, and ii) FSLE<0.05 $d^{-2}$ indicating the absence of significant structures (non-LCS) with a value equal to zero [18; Table S1]. OW and ADT were treated as continuous variables. The homogeneity of dispersions within each factor in the RDA was tested using the betadisper function from also in vegan [19]. Inorganic nutrient concentrations and $N_2$ fixation rates were only measured in zone 3 and thus not included in the RDA. The relation of these parameters with diazotroph abundances in zone 3 was then checked by Spearman rank correlations.

The significance of interactions between the abundance of each diazotroph group and the fine scale structure factors was tested by multivariate analysis of variance (ANOVA). To that end, fine scale variables were categorized (Table S1; Fig. S1). The OW parameter was categorized as “core” or “edge”, where core regions are under the influence of an eddy (OW<0) and edge regions represent a frontier where shear dominates (OW>0) [20]. FSLE was categorized as LCS or non-LCS as described above (Table S1). ADT was split into three levels (“low”, “medium” and “high”) according to the distribution of data across the three sampled zones (Fig. S1).

#### 2. Supplementary Tables

**Table S1:** Categorical classification of the physical parameters Okubo-Weiss (OW), finite size Lyapunov exponents (FSLE), and absolute dynamic topography (ADT) as used for statistical tests.

| Station | Zone | OW ($d^{-1}$) | OW categorical | FSLE ($d^{-2}$) | FSLE categorical | ADT (m) | ADT categorical |
|---|---|---|---|---|---|---|---|
| 153 | zone1 | -0.46 | core | 0 | non-LCS | 1.19 | high |
| 154 | zone1 | -0.49 | core | 0 | non-LCS | 1.17 | mid |
| 155 | zone1 | -0.38 | core | 0 | non-LCS | 1.17 | mid |
| 156 | zone1 | -0.12 | core | 0 | non-LCS | 1.17 | mid |
| 157 | zone1 | -0.11 | core | 0 | non-LCS | 1.14 | mid |
| 158 | zone1 | 0.42 | edge | 0 | non-LCS | 1.14 | mid |
| 159 | zone1 | 0.5 | edge | 0.39 | LCS | 1.14 | mid |
| 160 | zone1 | 0.41 | edge | 0.34 | LCS | 1.14 | mid |
| 161 | zone1 | 0.33 | edge | 0 | non-LCS | 1.13 | mid |
| 162 | zone1 | 0.2 | edge | 0.19 | LCS | 1.13 | mid |
| 163 | zone1 | -0.14 | core | 0 | non-LCS | 1.13 | mid |
| 164 | zone1 | -0.2 | core | 0 | non-LCS | 1.13 | mid |
| 165 | zone1 | -0.06 | core | 0 | non-LCS | 1.13 | mid |
| 166 | zone1 | 0.19 | edge | 0 | non-LCS | 1.13 | mid |
| 167 | zone1 | 0.18 | edge | 0 | non-LCS | 1.14 | mid |
| 168 | zone1 | 0.18 | edge | 0 | non-LCS | 1.14 | mid |
| 169 | zone1 | 0.26 | edge | 0.33 | LCS | 1.14 | mid |
| 170 | zone1 | 0.36 | edge | 0.17 | LCS | 1.14 | mid |
| 171 | zone2 | 0.08 | edge | 0 | non-LCS | 1.14 | mid |
| 172 | zone2 | 0.11 | edge | 0 | non-LCS | 1.14 | mid |
| 173 | zone2 | 0.3 | edge | 0 | non-LCS | 1.14 | mid |
| 174 | zone2 | 0.54 | edge | 0.12 | LCS | 1.1 | low |
| 175 | zone2 | 0.09 | edge | 0.39 | LCS | 1.13 | mid |
| 176 | zone2 | -0.31 | core | 0 | non-LCS | 1.1 | low |

| | | | | | | | |
|---|---|---|---|---|---|---|---|
| 177 | zone2 | -0.69 | core | 0.18 | LCS | 1.1 | low |
| 178 | zone2 | -0.63 | core | 0.18 | LCS | 1.07 | low |
| 179 | zone2 | -0.17 | core | 0.18 | LCS | 1.06 | low |
| 180 | zone2 | -0.18 | core | 0.17 | LCS | 1.06 | low |
| 181 | zone2 | -0.04 | core | 0 | non-LCS | 1.06 | low |
| 182 | zone2 | -0.22 | core | 0 | non-LCS | 1.08 | low |
| 183 | zone2 | 0.11 | edge | 0 | non-LCS | 1.16 | mid |
| 184 | zone2 | 0.17 | edge | 0.23 | LCS | 1.16 | mid |
| 185 | zone2 | 0.18 | edge | 0.2 | LCS | 1.21 | high |
| 186 | zone2 | -0.09 | core | 0.14 | LCS | 1.25 | high |
| 187 | zone2 | -0.55 | core | 0.13 | LCS | 1.25 | high |
| 188 | zone2 | -0.7 | core | 0 | non-LCS | 1.28 | high |
| 189 | zone2 | -0.05 | core | 0.16 | LCS | 1.27 | high |
| 190 | zone2 | 0.21 | edge | 0.14 | LCS | 1.25 | high |
| 191 | zone2 | 0.4 | edge | 0 | non-LCS | 1.23 | high |
| 192 | zone2 | 0.35 | edge | 0 | non-LCS | 1.23 | high |
| 193 | zone2 | 0.3 | edge | 0.11 | LCS | 1.22 | high |
| 194 | zone2 | 0.26 | edge | 0.14 | LCS | 1.21 | high |
| 195 | zone2 | 0.09 | edge | 0 | non-LCS | 1.21 | high |
| 196 | zone2 | 0.04 | edge | 0 | non-LCS | 1.22 | high |
| 197 | zone2 | -0.08 | core | 0.16 | LCS | 1.22 | high |
| 198 | zone2 | -0.16 | core | 0.2 | LCS | 1.22 | high |
| 199 | zone2 | -0.06 | core | 0.2 | LCS | 1.22 | high |
| 200 | zone2 | -0.01 | core | 0 | non-LCS | 1.22 | high |
| 201 | zone3 | -0.04 | core | 0 | non-LCS | 1.17 | mid |
| 202 | zone3 | -0.66 | core | 0.23 | LCS | 1.17 | mid |
| 203 | zone3 | -0.77 | core | 0 | non-LCS | 1.14 | mid |
| 204 | zone3 | -0.93 | core | 0 | non-LCS | 1.14 | mid |
| 205 | zone3 | -1.1 | core | 0 | non-LCS | 1.14 | mid |
| 206 | zone3 | -1.15 | core | 0 | non-LCS | 1.14 | mid |
| 207 | zone3 | -1.08 | core | 0 | non-LCS | 1.13 | mid |
| 208 | zone3 | -0.82 | core | 0 | non-LCS | 1.13 | mid |

| | | | | | | | |
|---|---|---|---|---|---|---|---|
| 209 | zone3 | -0.79 | core | 0 | non-LCS | 1.13 | mid |
| 210 | zone3 | -0.65 | core | 0 | non-LCS | 1.14 | mid |
| 211 | zone3 | 0.08 | edge | 0 | non-LCS | 1.14 | mid |
| 212 | zone3 | 0.16 | edge | 0 | non-LCS | 1.14 | mid |
| 213 | zone3 | 0.25 | edge | 0 | non-LCS | 1.17 | mid |
| 214 | zone3 | 0.79 | edge | 0.19 | LCS | 1.17 | mid |
| 215 | zone3 | 0.86 | edge | 0.43 | LCS | 1.17 | mid |
| 216 | zone3 | 0.82 | edge | 0.19 | LCS | 1.19 | high |
| 217 | zone3 | 0.72 | edge | 0 | non-LCS | 1.19 | high |
| 218 | zone3 | 0.63 | edge | 0 | non-LCS | 1.19 | high |
| 219 | zone3 | 0.56 | edge | 0.27 | LCS | 1.19 | high |
| 220 | zone3 | 0.12 | edge | 0 | non-LCS | 1.18 | high |
| 221 | zone3 | 0.01 | edge | 0 | non-LCS | 1.18 | high |
| 222 | zone3 | -0.07 | core | 0.13 | LCS | 1.18 | high |
| 223 | zone3 | -0.06 | core | 0 | non-LCS | 1.17 | mid |
| 224 | zone3 | 0.21 | edge | 0 | non-LCS | 1.19 | high |
| 225 | zone3 | 0.11 | edge | 0 | non-LCS | 1.19 | high |
| 226 | zone3 | 0.19 | edge | 0.13 | LCS | 1.18 | mid |
| 227 | zone3 | -0.08 | core | 0 | non-LCS | 1.22 | high |
| 228 | zone3 | -0.02 | core | 0 | non-LCS | 1.19 | high |
| 229 | zone3 | -0.01 | core | 0.48 | LCS | 1.15 | mid |
| 230 | zone3 | -0.1 | core | 0 | non-LCS | 1.15 | mid |
| 231 | zone3 | -0.47 | core | 0.17 | LCS | 1.15 | mid |
| 232 | zone3 | -0.65 | core | 0 | non-LCS | 1.12 | low |
| 233 | zone3 | -0.87 | core | 0 | non-LCS | 1.11 | low |
| 234 | zone3 | -0.8 | core | 0 | non-LCS | 1.11 | low |
| 235 | zone3 | -0.68 | core | 0 | non-LCS | 1.11 | low |
| 236 | zone3 | -0.39 | core | 0.12 | LCS | 1.13 | mid |
| 237 | zone3 | -0.35 | core | 0.11 | LCS | 1.15 | mid |
| 238 | zone3 | 0.15 | edge | 0.2 | LCS | 1.15 | mid |
| 239 | zone3 | 0.13 | edge | 0.13 | LCS | 1.17 | mid |
| 240 | zone3 | 0.26 | edge | 0 | non-LCS | 1.17 | mid |

| | | | | | | | |
|---|---|---|---|---|---|---|---|
| 241 | zone3 | 0.3 | edge | 0.11 | LCS | 1.17 | mid |
| 242 | zone3 | 0.11 | edge | 0 | non-LCS | 1.16 | mid |
| 243 | zone3 | 0.08 | edge | 0 | non-LCS | 1.16 | mid |
| 244 | zone3 | 0.02 | edge | 0 | non-LCS | 1.16 | mid |
| 245 | zone3 | 0.06 | edge | 0 | non-LCS | 1.15 | mid |
| 246 | zone3 | 0.09 | edge | 0 | non-LCS | 1.15 | mid |
| 247 | zone3 | 0.12 | edge | 0 | non-LCS | 1.15 | mid |
| 248 | zone3 | 0.09 | edge | 0.19 | LCS | 1.13 | mid |
| 249 | zone3 | 0.08 | edge | 0.22 | LCS | 1.16 | mid |
| 250 | zone3 | 0.06 | edge | 0.23 | LCS | 1.16 | mid |
| 251 | zone3 | 0.06 | edge | 0.23 | LCS | 1.16 | mid |
| 252 | zone3 | 0.11 | edge | 0.21 | LCS | 1.16 | mid |
| 253 | zone3 | 0.14 | edge | 0.21 | LCS | 1.16 | mid |
| 254 | zone3 | 0.16 | edge | 0.23 | LCS | 1.16 | mid |
| 255 | zone3 | 0.16 | edge | 0.23 | LCS | 1.17 | mid |
| 256 | zone3 | 0.21 | edge | 0.12 | LCS | 1.17 | mid |
| 257 | zone3 | 0.17 | edge | 0 | non-LCS | 1.19 | high |
| 259 | zone3 | 0.06 | edge | 0 | non-LCS | 1.2 | high |
| 260 | zone3 | 0.03 | edge | 0.11 | LCS | 1.2 | high |
| 261 | zone3 | 0.01 | edge | 0.11 | LCS | 1.22 | high |
| 262 | zone3 | 0.03 | edge | 0 | non-LCS | 1.22 | high |
| 263 | zone3 | 0.08 | edge | 0 | non-LCS | 1.22 | high |
| 264 | zone3 | 0.09 | edge | 0 | non-LCS | 1.24 | high |
| 265 | zone3 | 0.28 | edge | 0 | non-LCS | 1.24 | high |
| 266 | zone3 | 0.24 | edge | 0 | non-LCS | 1.26 | high |
| 267 | zone3 | 0.17 | edge | 0 | non-LCS | 1.24 | high |
| 268 | zone3 | 0.13 | edge | 0 | non-LCS | 1.23 | high |
| 269 | zone3 | 0.01 | edge | 0 | non-LCS | 1.22 | high |
| 270 | zone3 | -0.03 | core | 0 | non-LCS | 1.22 | high |

**Table S2:** The impact of physical parameters on diazotroph abundance based on one-way ANOVA statistical analyses. Significant results ($p \leq 0.05$) are depicted with an asterisk.

| Diazotroph group | Physical parameter | F value | p-value |
|---|---|---|---|
| *Trichodesmium* | OW | 1.691 | 0.196 |
| | FSLE | 0.005 | 0.944 |
| | ADT | 11.03 | *4.61 $x10^{-5}$ |
| UCYN-A | OW | 0.68 | 0.998 |
| | FSLE | 3.731 | *0.050 |
| | ADT | 2.693 | 0.072 |
| UCYN-B | OW | 1.601 | 0.209 |
| | FSLE | 0.6 | 0.441 |
| | ADT | 1.867 | 0.160 |
| UCYN-C | OW | 3.146 | 0.078 |
| | FSLE | 5.248 | *0.0239 |
| | ADT | 5.2 | *0.007 |
| Gamma A | OW | 2.709 | 0.103 |
| | FSLE | 0.452 | 0.503 |
| | ADT | 10.27 | *7.95 $x10^{-5}$ |

#### 3. Supplementary Figures

**Fig. S1:** Histogram of absolute dynamic topography (ADT) values and divisions into low, mid and high levels. The differences between each pair of groups is statistically significant as confirmed by t-tests (low vs mid $p = 8.69 \times 10^{-7}$, mid vs high $p = 2.2 \times 10^{-16}$ and low vs high $p = 5.06 \times 10^{-13}$).

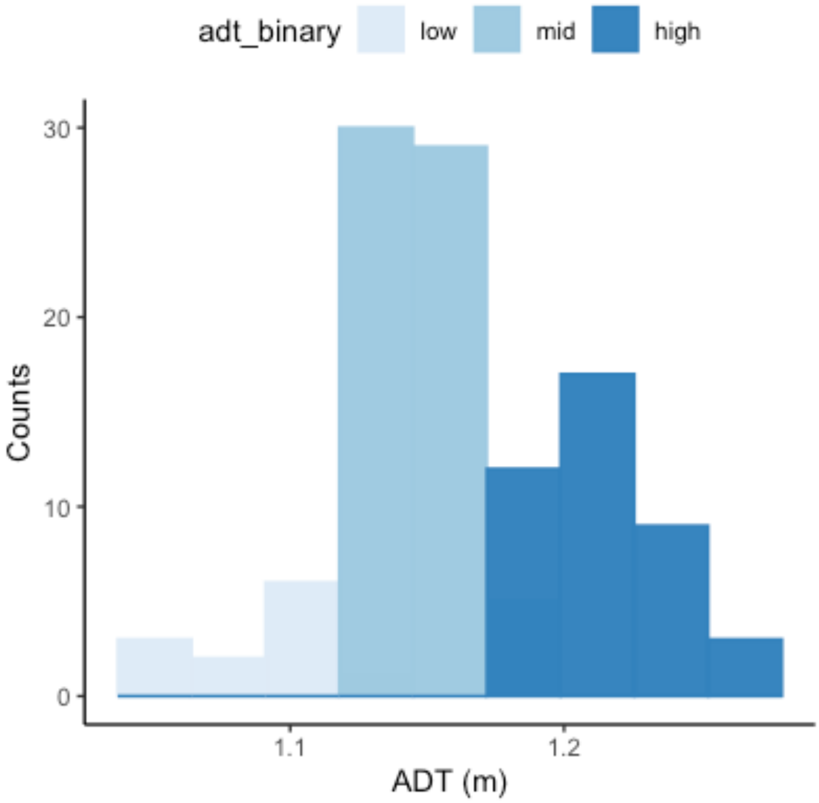

**Fig. S2:** Multivariate redundance analysis (RDA) biplot depicting the variance explained by the fine scale parameters and qPCR data. The RDA1 and RDA2 axes explained 56.13 and 36.25% of the variance, respectively.

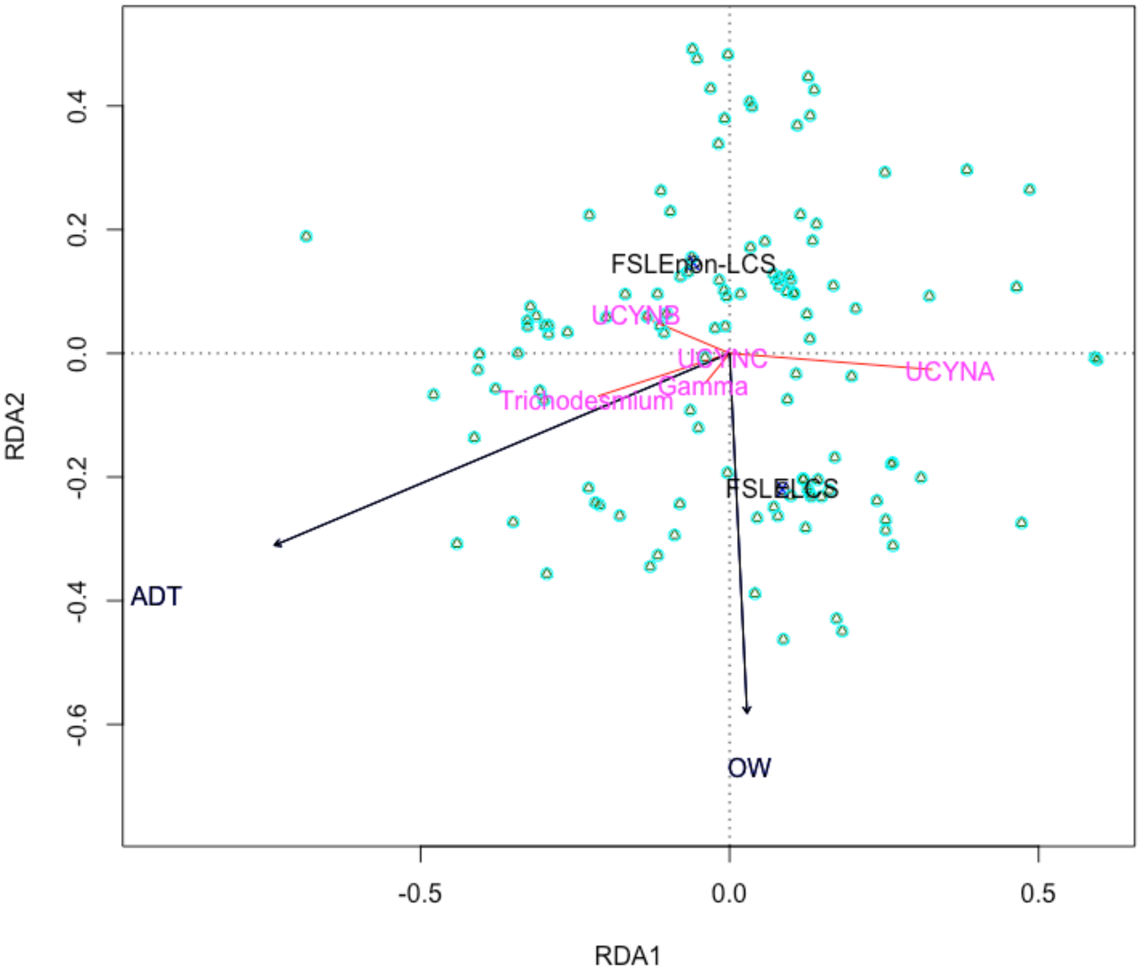

**Fig. S3: Pearson correlations between biological and environmental variables.**

Biological variables (*nifH* gene counts of *Trichodesmium,* UCYN-A, UCYN-B, UCYN-C and Gamma) and environmental variables (temperature, salinity and fluorescence). Statistically significant ($p \leq 0.05$) positive and negative correlations are shown in red and blue circles, respectively. The size of the circles indicates the magnitude of the p value, while the color tone indicates the Pearson correlation coefficient value as depicted in the color scale.

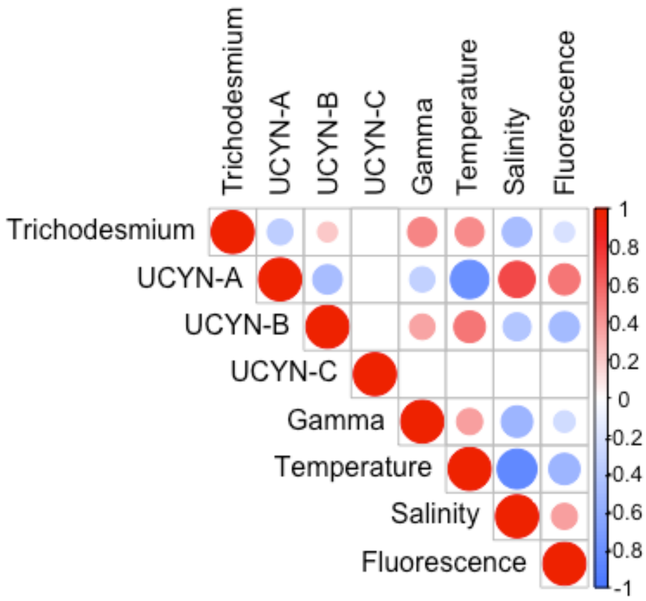

**Fig. S4: Underway temperature, salinity and fluorescence in zones 1, 2 and 3.** Data were retrieved for each zone on 2nd, 4th and 22nd November 2019 for zones 1, 2 and 3, respectively.

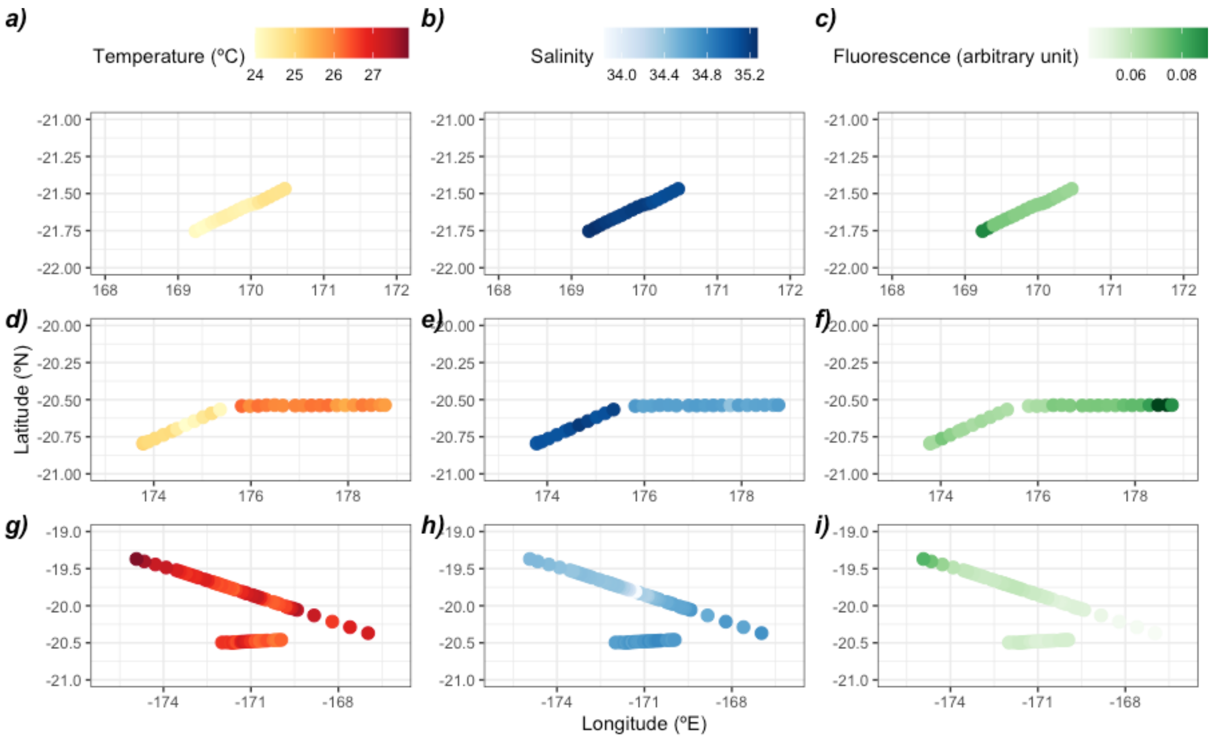

**Fig. S5: Distribution of inorganic nutrients (nitrate and phosphate) and $N_2$ fixation rates in zone 3.** Data were retrieved for each zone on 2nd, 4th and 22nd November 2019 for zones 1, 2 and 3, respectively. Grey dots depict locations where samples for diazotroph abundance are available (see Fig. 1 and Fig. S3), but inorganic nutrient concentration and $N_2$ fixation rate data are not available.

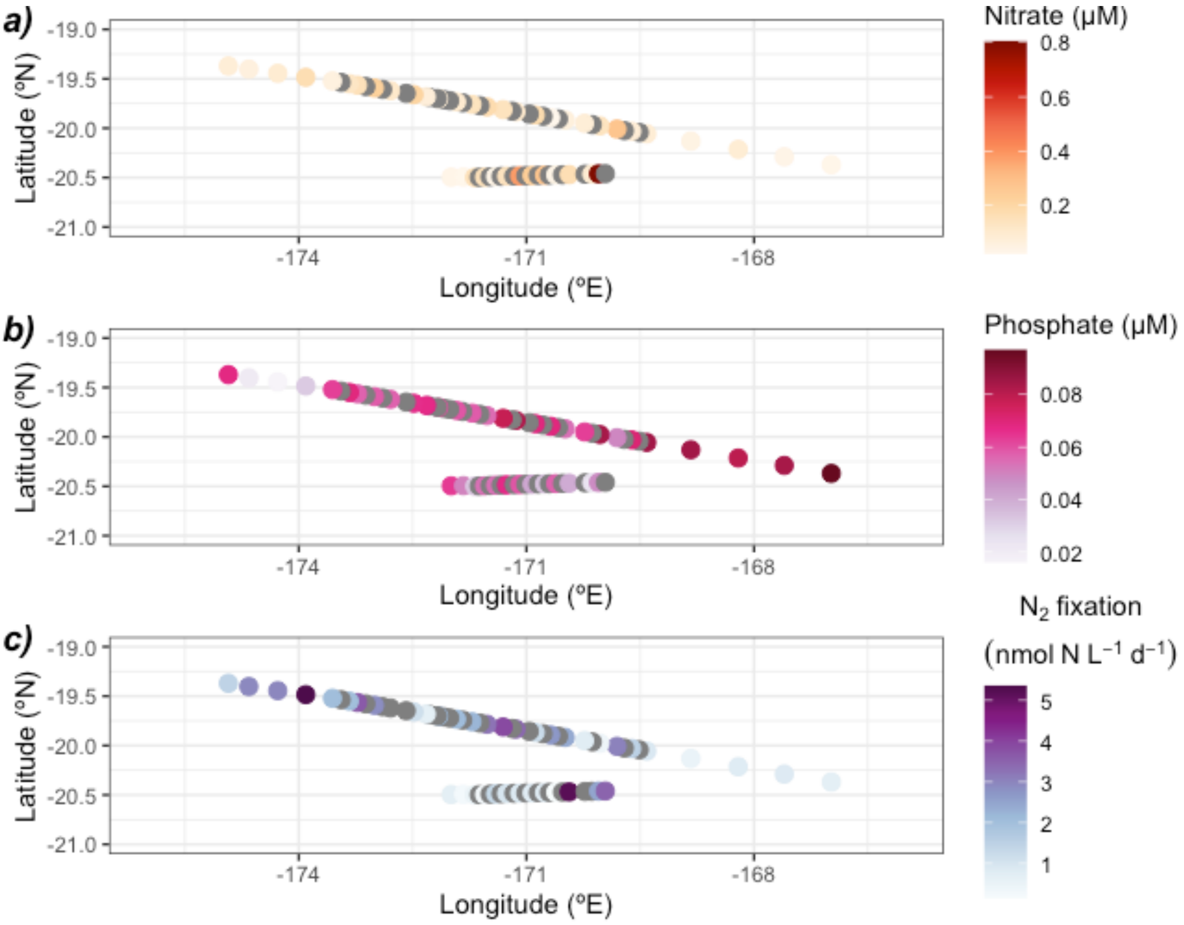
